# A class I hydrophobin in *Trichoderma virens* influences plant-microbe interactions through enhancement of enzyme activity and MAMP recognition

**DOI:** 10.1101/2021.01.07.425738

**Authors:** James T. Taylor, Inna Krieger, Frankie K. Crutcher, Pierce Jamieson, Benjamin A. Horwitz, Michael V. Kolomiets, Charles M. Kenerley

## Abstract

The filamentous fungus, *Trichoderma virens*, is a well-known mycoparasitic plant symbiont, valued for its biocontrol capabilities. *T. virens* initiates a symbiotic relationship with a plant host through the colonization of its roots. To achieve colonization, the fungus must communicate with the host and evade its innate defenses. Hydrophobins from *Trichoderma spp.* have previously been demonstrated to be involved in colonization of host roots. In this study, the class I hydrophobin, HFB9A from *T. virens* was characterized for a potential role in root colonization. Δhfb9a gene deletion mutants colonized less than the wild-type strain, were unable to induce systemic resistance against *Colletotrichum graminicola*, and showed a reduction in the activity of its cell wall degrading enzymes. The purified HFB9A protein was able to complement the enzyme activity of mutant culture filtrates as well as enhance the activity of commercially sourced cellulase. When exogenously applied to Arabidopsis plants, HFB9A protein induced phosphorylation of AtMAPK3/6, suggesting that it functions as a microbe-associated molecular pattern.

## Introduction

The filamentous plant symbiotic fungus *Trichoderma virens* is recognized for its ability to colonize plant roots and provide benefits to its hosts through the induction of systemic resistance, protection against fungal root pathogens, and growth promotion (Howell, 1987; Pieterse *et al.*, 2014; Saldajeno *et al.*, 2014). During the colonization process, a large number of secreted fungal proteins are involved in the subroutines of evading the plant defenses, initially penetrating roots, and fungal growth within the root system by hyphal expansion (Djonović*et al.*, 2006a; Crutcher *et al.*, 2015; Lamdan *et al.*, 2015). In addition to fungal proteins that are secreted into intercellular spaces, others localize to the outer cell wall of the fungus. Here, they can serve a wide variety of functions such as receptors for specific stimuli, protect against antimicrobial compounds, and aid in physical interactions including attachment to surfaces (Zampieri *et al.*, 2010; Bignell, 2012; Kim *et al.*, 2016; Correia *et al.*, 2017). Much effort has been extended to discover and understand proteins involved in *Trichoderma-plant* interactions, with a major emphasis on small, secreted cysteine-rich proteins hypothesized to function as effectors (Lamdan *et al.*, 2015; Morán-Diez *et al.*, 2015; Guzmán-Guzmán *et al.*, 2017; Ramírez-Valdespino *et al.*, 2019). The best-known example of this type of protein is SM1, which belongs to the cerato-platanin family and is required for induced systemic resistance (ISR) mediated by *T. virens* (Djonović*et al.*, 2006a; Djonović *et al.*, 2007). In sharp contrast, the role of other secreted protein families in *Trichoderma-plant* interactions are much less understood. Of these families, hydrophobin proteins are of particular interest due to their diverse suite of functions.

Hydrophobins are small cysteine-rich, secreted proteins that self-assemble at hydrophilic/hydrophobic interfaces (Wösten, 2001) and are unique to fungi. They contain a conserved motif of cysteine residues, which they may act as effectors involved in plant-fungal interactions, similarly to other cysteine-rich proteins (Ruocco *et al.*, 2015; Guzmán-Guzmán *et al.*, 2017). As secreted proteins, some hydrophobins cover the surface of spores of fungal pathogens, helping to evade host defenses (Bayry *et al.*, 2012). Others aid fungal morphogenesis by enabling hyphae to penetrate air/water interfaces (Wösten and de Vocht, 2000). The unique properties of hydrophobins are suited for a variety of applications in industrial and scientific techniques. The fusion of a hydrophobin to a protein of interest can significantly boost the yield of the purified protein (Joensuu *et al.*, 2010; Mustalahti *et al.*, 2013). Hydrophobins are also industrially used as emulsifiers and agents to alter surface characteristics of substrates (Bayry *et al.*, 2012).

Hydrophobins are currently organized into two classes (I and II) based on solubility, the spacing between cysteines, and hydrophobicity patterns in the amino acid sequence (Wösten, 2001). Class II hydrophobins are more soluble than class I and have more conserved spacing between cysteines, whereas class I hydrophobins are very insoluble, requiring harsh acids to dissolve, and can have highly variable cysteine spacing (Wösten and De Vocht, 2000; Wösten, 2001; Bayry *et al.*, 2012). Class I hydrophobins typically localize to the outer cell wall of fungal hyphae and/or spores where they form monolayers or self-assemble into amyloid-like fibrils. This differs from class II hydrophobins, which tend to be freely secreted into the environment. A limited number of hydrophobins from *Trichoderma* species have been functionally characterized. A class I hydrophobin, TASHYD1 from *T. asperellum*, was found to aid in the attachment of conidia and hyphae to roots for more efficient colonization of cucumber plants (Viterbo and Chet, 2006). A class II hydrophobin, HYTLO1 from *T. longibrachiatum*, was shown to exhibit direct antifungal effects and induce systemic resistance when applied to plant leaves (Ruocco *et al.*, 2015). Additionally, a class II hydrophobin from *T. virens* was demonstrated to have a role in root colonization and mycoparasitism activity (Guzmán-Guzmán *et al.*, 2017). Class I hydrophobins have been identified in *T. virens*, but are fewer in number (three) than class II hydrophobins (eight), and no distinctive role in plant interactions has been demonstrated (Seidl-Seiboth *et al.*, 2011). One class I hydrophobin, HFB9A from *T. virens*, has been described that shares significant homology with TASHYD1 (Viterbo and Chet, 2006). Based on this homology and the typical characteristics of hydrophobins, we hypothesized that HFB9A has a role in *T. virens*-plant interactions. In this study, we demonstrate the function of the hfb9a gene in root colonization and induction of systemic resistance in maize as well as the enhancement of enzyme activity on cell wall components.

## Materials and Methods

### Bioinformatic analysis

The protein sequences of selected proteins were subjected to a BLAST search of the NCBI database for homologs. The resulting matches were aligned using CLUSTAL Omega software. Additionally, the DNA sequence of the promoter and terminator regions of the hydrophobin were queried through the Joint Genome Institute BLAST (https://mycocosm.jgi.doe.gov/pages/blast-query.jsf?db=TriviGv29_8_2) search of the *T. virens Gv29-8* genome.

### Strains and conditions

The root pathogens, *Pythium ultimum* and *Rhizoctonia solani*, and the wild-type strain of *T. virens* [Gv29-8] were maintained on potato dextrose agar (PDA, BD Difco™) at 27 °C. The maize foliar pathogen *Colletotrichum graminicola* was maintained under an 14:10 light:dark light regime at room temperature on PDA plates for sporulation. Chlamydospores of *T. virens* were harvested from 14-day old cultures of *T. virens* grown in Fernbach flasks containing 1 L of molasses medium (30g molasses and 5g yeast extract per liter of water) by vacuum filtration and dried overnight. The dried chlamydospore mats were ground in a Wiley mill with a #60 sieve. *Zea mays*(Silver queen hybrid, Burpee) were grown in plastic cone containers in Metromix soilless medium or in a hydroponic system (Lamdan *et al.*, 2015). The hydroponic system consisted of mason jars (500ml, wide mouth) with a shaker clamp placed inside. The jar was filled to the top of the clamp (~220ml) with 0.5x Murashige-Skoog basal medium containing Gamborg’s vitamins and supplemented with 0.5% sucrose. The unit was covered with a glass petri dish bottom and autoclaved. Plastic mesh (7 holes/linear inch) previously cut into discs to fit within the jars was autoclaved separately. After sterilization, the mesh discs were placed on top of the clamps, and pregerminated seeds with roots approximately 2 cm long were threaded through the mesh to contact the growth medium. The glass petri dish bottoms were replaced with sterile plastic petri dish bottoms, as they ensure a tighter fit. All plants were grown under lamps (Sun Blaze T5) with 6500K and 3000K lights at room temperature under a 14:10 light:dark regime.

### RNA isolation and Expression assays

Total RNA was extracted from cultures of *T. virens* grown in potato dextrose broth (PDB, BD Difco™) that were inoculated with approximately 3×10^9^ conidia. The fungal biomass was collected every 24 hr over the course of 7 days. To determine the expression of *hfb9a* in plant-fungal interactions, *T. virens* was grown in a hydroponic system in the presence of maize roots, and fungal tissue samples were collected at 6, 30, and 54 hr. All samples were extracted using the Direct-zol RNA miniprep kit (Zymo Research, USA) following manufacturer’s instructions. Extracted RNA was converted to cDNA using the high capacity cDNA reverse transcription kit (Applied Biosystems) according to manufacturer’s instructions. The resulting cDNA was analyzed by RT-PCR using gene specific primers and primers amplifying Histone H3 as a loading control (Supplementary Table 1). Raw read counts were obtained from previously performed RNA-seq transcriptomic studies (Taylor *et al., unpublished;* Malinich *et al.*, 2019). The reads were normalized using the TMM algorithm in EdgeR. The normalized reads were then queried for those corresponding to *hfb9a* and graphed using seaborn and matplotlib packages in python.

### Deletion of hfb9a

The hfb9a gene was targeted for deletion through homologous recombination using a vector generated via the OSCAR method (Paz *et al.*, 2011) modified for use in *T. virens.* Primers were designed to amplify approximately 1kb of upstream and downstream regions flanking the open reading frame of the gene and contained appropriate Gateway sites for recombination (Supplementary Table 1). The flanks amplified from *T. virens* genomic DNA were purified by adding 90 μl of combined PCR product, 270 μl TE buffer, and 180 μl 30% PEG 8000/30mM MgCl2 to a microfuge tube. This solution was vortexed thoroughly, and then centrifuged for 15 min at maximum speed in the table-top centrifuge. The pellet was resuspended in 15 μl of sterile water. A 5 μl clonase reaction was performed using 20 ng of combined flanks, 60 ng pA-Hyg-OSCAR, 60 ng pOSCAR, and 1 μl of BP clonase (Invitrogen) with incubation in a thermocycler overnight at 25 °C. The reaction was stopped the next morning by adding 0.5 μl of proteinase K and incubating at 37 °C for 10 min. The entire reaction was used to transform *E. coli* DH5α cells and positive clones were screened as described in Paz et al (2011).

The resulting vector (pHFB9a) was electroporated into *A. tumefaciens* AGL1 and confirmed through colony PCR using primers to amplify both flanks and the hygromycin resistance gene (hph) (Supplementary Table 1). Overnight cultures of AGL1 containing the vector were pelleted and resuspended to an OD600 of 0.15 in induction medium (M9 minimal medium; 100 ml 5x M9 salts, 3.9g MES, 0.45g glucose, 0.25 ml glycerol, to 500 ml with H2O, pH 5.3) with and without 200 μM acetosyringone and allowed to incubate for 6 hr at 27 °C. Conidia of *T. virens* were collected from 4-day old PDA plates, diluted to 5×10^5^ conidia/ml, and mixed with bacteria in a 1:1 ratio. Several sterile cellophane squares (~1cm^2^) were placed on co-cultivation plates (M9 minimal medium with 500 μM acetosyringone and 1.5% agar) and 20 μl of the mixed conidia/bacteria solution were placed on each square. The plates were allowed to incubate for 60 hr before transferring the cellophane to PDA selection plates containing hygromycin, tetracycline, and chloramphenicol. Positive transformants were transferred to 2 ml PDA slants containing the same antibiotics. Once cultures began to sporulate, they were successively transferred to PDA + antibiotics, PDA, and back to PDA + antibiotics to ensure stability of the integration. Stable transformants were grown in PDB for 2 days and genomic DNA extracted for PCR analysis. Primers specific to the ORF of the gene and primers outside the 5’ flank and inside of hph were used to confirm deletion.

### Phenotypic analysis of mutants

Two mutants and the wild-type were assayed for differences in general morphology and radial growth rate by plating a 3 mm radius plug of actively growing fungus on four PDA plates each and measured every day for four days. The strains were also tested for differences in mycoparasitic ability in confrontation with *P. ultimum*. Plugs of each fungus were placed on opposite sides of a PDA plate 1 cm away from the edge of the plate and allowed to grow toward each other for seven days. The length of the growth front of wild-type or mutants was measured from the plug and recorded for comparison. Each experiment was repeated twice with four independent plates per experiment. Biocontrol activity was measured as in Djonović et al. using *R. solani* as the pathogen rather than *P. ultimum* (Djonović*et al.*, 2006b). Contact angle measurement was performed as in Crutcher et al. (Crutcher *et al.*, 2015).

### Oxidative stress assay

Oxidative stress tolerance was measured by growth on VMS agar plates containing 10 μM sodium menadione bisulfite. Agar plugs (3 mm radius) of each strain were placed in the middle of the plates. Radial growth was measured every 24 hr for three days.

### Root colonization and Induced Systemic Resistance assays

Maize seedlings were grown in a hydroponic system as described previously. Once the roots had reached sufficient length (approx. 3-5 cm), 1 g of tissue from wild-type or mutant *T. virens* strains was placed in the liquid growth medium and gently stirred to distribute. The seedlings and fungal biomass were incubated for 3 days shaking at 50 rpm. The roots were then harvested and thoroughly rinsed in tap water. The collected roots were ground in liquid nitrogen and genomic DNA was extracted using the same protocol described above. The samples were analyzed via the ΔΔCt method of qPCR with actin and phenylalanine ammonia lyase primers to determine the ratio of fungal to maize DNA, respectively (Crutcher et al. 2013). The maize samples were treated as the endogenous control and the WT: Maize DNA ratio was normalized to one relative abundance unit and used as the basis of comparison. Mutants were investigated for changes in ISR activity against the foliar pathogen *C. graminicola* following the protocol of Djonović et al. (Djonović *et al.*, 2007) using Silver Queen hybrid plants instead of the B73 inbred line. The area of individual lesions was measured using ImageJ (Schneider *et al.*, 2012). *C. graminicola* was utilized due to its status as a top maize pathogen and the consistency of the lesions that it causes. The shoot height of treated plants was measured with a meter stick after removing the plants from the plastic cone containers starting at the seed and ending at the longest leaf tip after straightening. After shoot measurements, the roots of the plants were cleaned under running water to remove attached soil and dried in an oven overnight. The combined dry weight of roots and shoots from each plant was recorded.

### Confocal microscopy and staining

Colonized sections of roots harvested from the hydroponic system after two days incubation with strains of *T. virens* were cleared by treatment with 10% KOH for 1 hour at 95C. The samples were equilibrated in PBS (pH 7.4) for 1 hour. The equilibrated samples were infiltrated in a solution of 5 mg/ml WGA-Alexa-fluor 488 and 10 mg/ml propidium iodide in PBS (pH 7.4) for 15 min under vacuum and destained in PBS for an additional 15 min. The stained samples were immediately visualized on an Olympus FV3000 confocal microscope.

### Enzymatic activity assay of cell wall degrading enzymes

Six replicates of 3 mm radius plugs of each strain were grown in 2 ml of VMS broth in a 24 well plate for 48 hr at 27°C. A 150 μl sample of broth from each well was added to a PCR tube along with 150 μl of Bradford reagent and incubated at room temperature for 5 min. To determine total protein concentrations, the absorbance of the samples was measured at 595 nm and compared to absorbance values of a standard BSA curve. Samples from each well were diluted to a total protein concentration of 10 μg/ml. These diluted samples were used for enzyme activity assays.

To measure cellulolytic activity, 40 μl of each sample were transferred to a well of a 96 well microplate and repeated for a total of 3 technical replicates per sample. A 60 μl aliquot of 50 mM sodium acetate buffer at a pH of 4.8 was added to the well along with 10 μl of 1% carboxymethyl cellulose. The plate was sealed with adhesive film and incubated in a thermocycler at 50°C for 60 min. A 50 μl aliquot of solution from each well was transferred to a new 96 well plate and 100 μl of dinitrosalicylic acid solution was added. The new plate was then incubated at 95°C for 5 min to allow color to develop, after which 40 μl of the developed solution was added to a 96 well plate, diluted with 160 μl H2O, and absorbance measured at 540nm in a microplate reader.

To measure chitinase activity, 20 μl of sample was added to a well of a flat bottom 96 well plate. To this, 80 μl of the same sodium acetate buffer used above and 5 μl of 0.5 mg/ml 4-methylumbelliferyl β-D-N, N’, N”-triacetylchitotrioside as a substrate was added. This mixture was incubated for 15 min at 40°C, and fluorescence was measured in a microplate reader.

### Protein expression and extraction

Primers were designed to amplify the 423 bp coding sequence from cDNA with *NdeI* and *HindIII* restriction sites prepended to the 5’ and 3’ primers, respectively (Supplementary Table 1). The amplicon and pET30b(+) were double digested with *Nde*I and *HindIII* for 30 min at 37°C. The digested products were cleaned with a QIAquick PCR cleanup kit (Qiagen, US) and ligated at a 3:1 amplicon:plasmid ratio with t4 DNA ligase (NEB, US) overnight at 16°C. A 5 μl aliquot of the ligase mix was transformed into DH5α competent cells via heat shock. Following a recovery period of one hour in SOC medium, 200 μl were plated on an LBA plate containing 50 μg/ml kanamycin and incubated overnight at 37°C. Resulting colonies were screened by PCR for amplification of the ORF of hfb9a and positive colonies were digested with *Nde*I and *Hind*III to determine insert size. Several vectors were then sequenced for confirmation of the correct insertion.

The confirmed vector was transformed into *E. coli* BL21(DE3) competent cells (Invitrogen, USA) via heat shock. Four resulting colonies were inoculated into 3 ml of LB broth containing 50 μg/ml kanamycin. The cultures were shaken at 225 rpm in a 37°C incubator until the OD600 reached approximately 0.6 (roughly 5 hr). Expression was induced by adding 0.75 mM IPTG to the cultures with further incubation at 37°C for 4 hr. The cultures were transferred to 2 ml Eppendorf tubes and centrifuged for 5 min at 4500xg. The resulting pellet was washed with 500 μl of phosphate buffered saline (PBS, pH 7), and subsequently resuspended in 500 μl of PBS. The cells were lysed by repeated freeze-thaw cycles with liquid nitrogen. A 100 μl sample of the solution was collected in a 1.5 ml Eppendorf tube to represent the combined soluble and insoluble fractions of the lysate. The remainder was centrifuged for 5 min at full speed in a tabletop centrifuge. A 100 μl sample was taken to represent the soluble fraction of the lysate. Each sample was diluted with gel loading buffer and placed in a boiling water bath for 5 min. 10 μl of each sample were loaded into a 15% SDS-PAGE gel and run at 35 mA for 1 hour.

To assess the presence of the protein of interest, a dot blot was performed using anti-His antibodies conjugated to alkaline phosphatase. For each sample, 3 μl of each protein extract and a positive control were dotted onto a nitrocellulose membrane. After drying, the membrane was blocked using 5% skim milk powder in TTBS for 30 min. The membrane was rinsed briefly with TTBS, and the dilute antibody solution was added and allowed to incubate for 1 hour. The antibody solution was drained off, and the membrane was thoroughly rinsed with TTBS. The membrane was developed by adding 10 ml of BCIP/NBT (Sigma Aldrich, USA) solution to the membrane and incubating for 20 min.

An overnight culture of the expressing strain of *E. coli* was used to start a 2 L culture. The culture was centrifuged for 50 min at 16,000 rpm with the pellet resuspended in lysis buffer (20 mM Tris pH 7.5, 100 mM NaCl) and 2 M urea. Then 25 μl of 1 M MgCl2 and 25 μl of DNaseI were added following resuspension, the solution was lysed using a French press, and centrifuged at 16,000 rpm for 50 min to obtain a pellet. The supernatant was decanted into a separate container for later testing. The pellet was washed twice by resuspending in lysis buffer + 2 M urea and spun at 16000 rpm for 20 min. The final pellet was resuspended in lysis buffer + 8 M urea. The suspension was spun once more at 16,000 rpm for 20 min. The supernatant containing solubilized inclusion bodies was passed through a HisTrap HP (5 ml column volume, GE, USA) chromatography column using a peristaltic pump. The column was then attached to an FPLC where the bound proteins could be refolded by passing lysis buffer containing a slow gradient (20 column volumes) from 8 M to 0 M urea with 5 mM reduced glutathione and 0.5 mM oxidized glutathione. Following refolding, the protein was eluted by 0-400 mM imidazole gradient. The fractions were analyzed by SDS-PAGE gel, as well as subjected to a dot blot with anti-his antibodies to confirm the presence of the recombinant protein.

### SM1 production determination

Cultures of each strain were grown in 1 L VMS shaken at 150 rpm and 27C for one week. The cultures were filtered through Whatman #4 filter paper and the filtrate collected. Proteins were precipitated from the filtrate with ammonium sulfate (~80% saturation) and collected by centrifugation. Levels of SM1 production were determined by immunoblotting as in Djonovic *et al.* 2006a.

### Arabidopsis MAPK phosphorylation assay

Arabidopsis seedlings were grown on plates of 0.5x Murashige-Skoog basal medium with Gamborg’s vitamins for ten days. Several seedlings were placed in wells of a 12 well plate with 500 ul of sterile water and incubated overnight. The seedlings were then treated with chitin, purified HFB9A, the protein suspension buffer, or purified SM1. The proteins were added to a concentration of 100 nM. After 15 or 30 min, the seedlings were collected, and flash frozen in liquid nitrogen. The frozen seedlings were ground in protein extraction buffer (Li *et al.*, 2015) and run on a 10% SDS-PAGE gel. The proteins were transferred and blotted with anti-pERK1/2 antibodies and detected by enhanced chemiluminescence.

### Statistical analysis

All data was analyzed for statistical significance using the ANOVA, Tukey’s HSD, and/or Kruskal-Wallis functions in R.

## Results

### The T. virens genome encodes two canonical type I hydrophobins

NCBI PSI-BLAST search using the protein sequence of HFB9A (Genbank Accession: EHK16816.1) revealed a characterized homolog in the *T. asperellum* genome, TASHYD1. A second similar hydrophobin was found in the *T. virens* genome that has not yet been characterized. With MEGA software, a phylogenetic tree was constructed using selected top hits of the NCBI PSI-BLAST, with HFB9A serving as the query sequence (Figure 1). The tree diverged into two main clades representing proteins that were more similar to HFB9A or HFB3A. The only characterized hydrophobin in the phylogeny was TASHYD1, which clustered with HFB9A, indicating that the two proteins may share similar characteristics and roles in the fungus. Additionally, PFAM database scanning revealed a hydrophobin domain with an N-terminal signal peptide and no other conserved domains in the amino acid sequence. The protein sequence of HFB9A was used for homology modeling of protein structure with the I-TASSER software (Yang *et al.*, 2014). The predicted structure of HFB9A (Figure 2) shared structural homology with human defensin and cell adhesion proteins and was predicted to bind a peptide as a ligand. The structure of TASHYD1 was modeled using the same software. Both HFB9A and TASHYD1 successfully model as similar to the solved structure of DEWA from *Aspergillus nidulans* (Morris *et al.*, 2013) showing the similar core of beta-sheets, with the unique composition of the surface residues most likely responsible for the specific functions.

**Figure 1.**
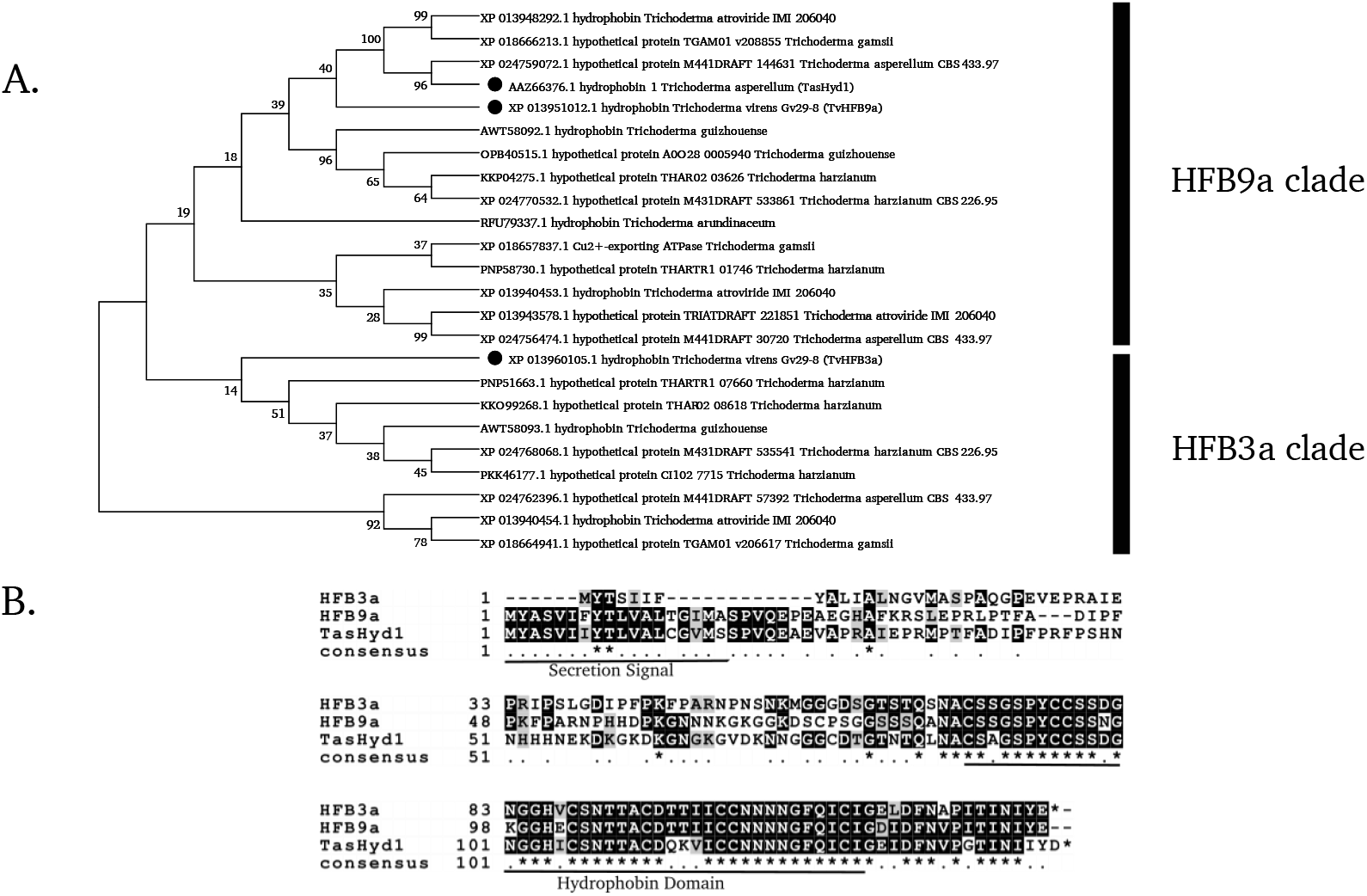
Phylogenetic comparison of hydrophobins from *Trichoderma spp.***A.**A phylogenetic tree of class I hydrophobins from *Trichoderma spp.* produced with MEGA software. **B.**A Clustal-Omega alignment of the amino acid sequences of two *T. virens* class I hydrophobins (HFB9a and HFB3a) and TASHYD1 from *T. asperellum.* The residues that make up the N-terminal secretion signal and hydrophobin core are labeled with an underline. Identical residues between all three sequences are labeled with an asterisk.

**Figure 2.**
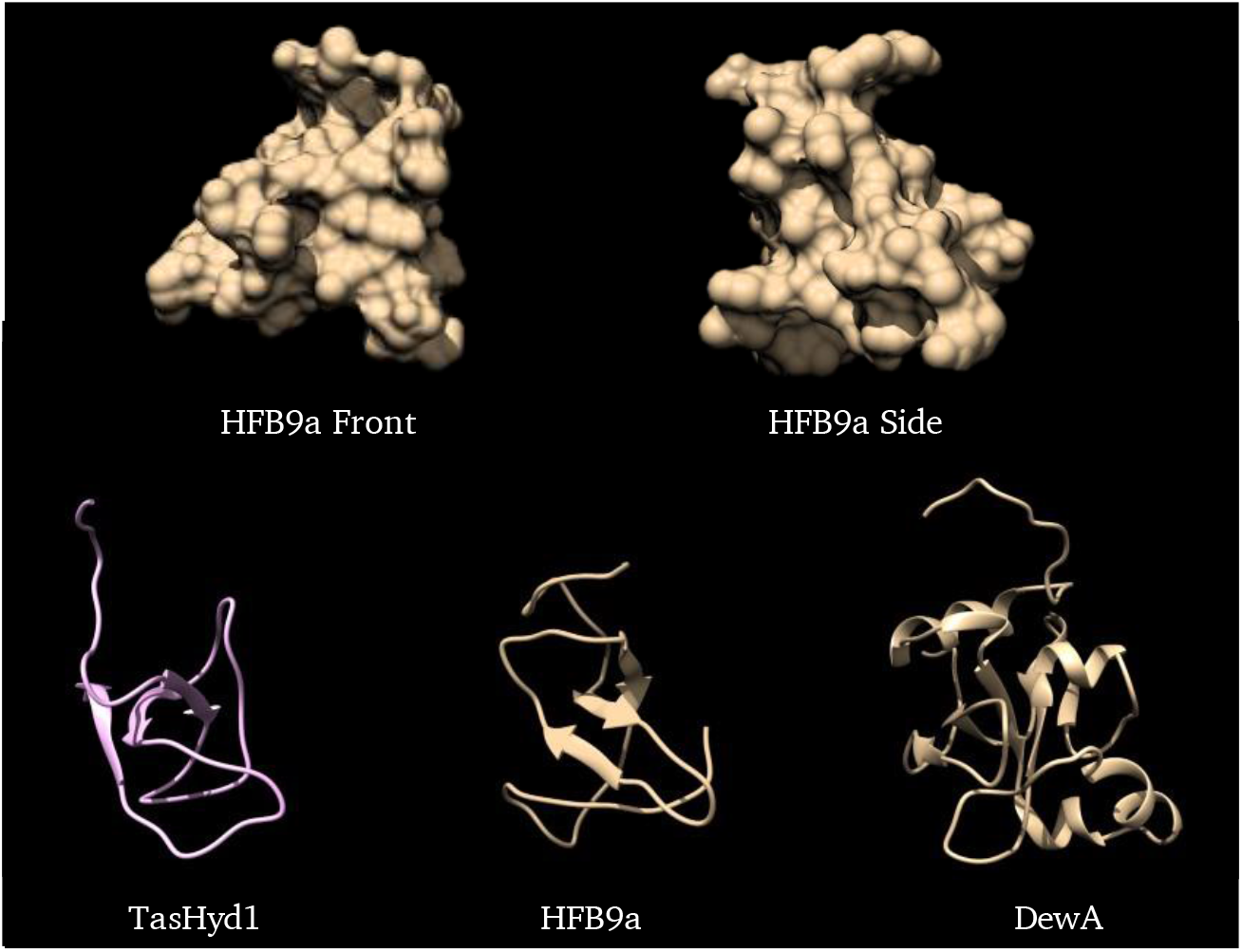
A comparison of the predicted protein structures of TASHYD1, HFB9a, and the solved structure of DEWA, a class I hydrophobin from *Aspergillus nidulans.* Both HFB9A and TASHYD1 successfully model as similar to the solved structure of DEWA from *Aspergillus nid-ulans* (Morris *et al.*, 2013) showing the similar core of beta-sheets, with the unique composition of the surface residues most likely responsible for the specific functions.

### hfb9a is induced during fungal association with maize roots

To develop an expression profile for *hfb9a*, cDNA generated from RNA of wild-type collected at predetermined time points either in the presence of living maize roots (6, 30, and 54 hpi) or in shaken culture of potato dextrose broth (PDB, collected every day for 7 days) was subjected to RT-PCR. Expression was observed only in samples from the hydroponic system collected 54 hr post inoculation, indicating expression of the gene between 30 and 54 hpi (Figure 3A). Expression of the gene from mycelial samples grown in PDB was not detected until 72 hpi but remained constant in the remainder of the samples. In the hydroponic system, attachment of the wild-type and each mutant to the root system of maize seedlings was recorded at approximately 6 hpi, with the entirety of the root system enveloped by fungus at 54 hpi. The initial expression results were further confirmed by whole transcriptome sequencing data (Figure 3B, 3C, Malinich *et al.*, 2019, Taylor *et al. unpublished).* In early time points (6-24 hpi, Figure 3B, 3C), the normalized transcript counts are near 0, while starting at 30 hr post inoculation transcript counts rapidly increase.

**Figure 3.**
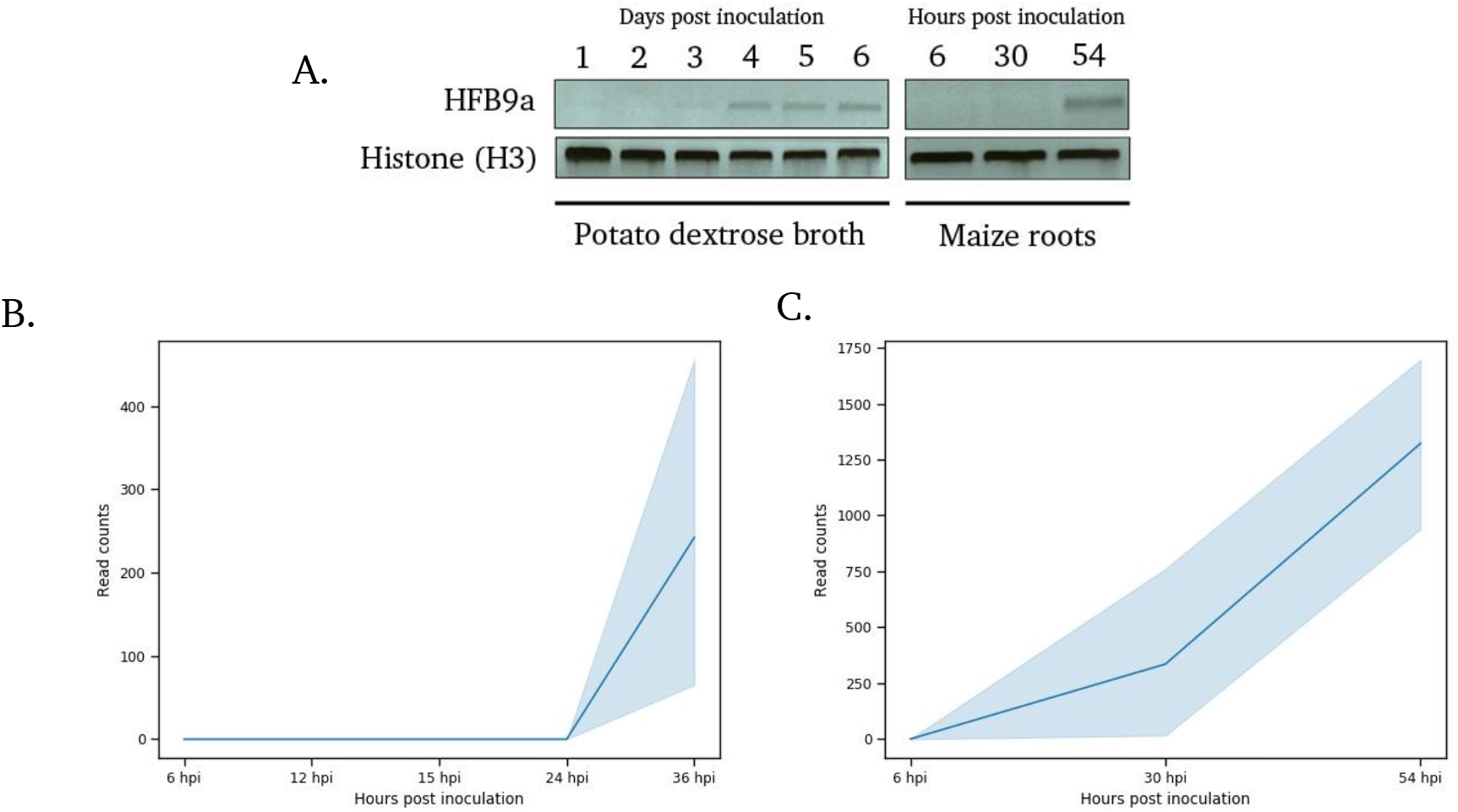
Expression profiling of *hfb9a* **A**. *hfb9a* expression in potato dextrose broth and in the presence of maize roots as measured by RT-PCR. Histone (H3) was used as a loading control. **B and C.**Normalized read counts from RNA-seq based transcriptomic datasets plotted across time (B: Taylor *et al.* unpublished, C: Malinich *et al.*, 2019). Raw read counts were obtained from the mentioned studies, then normalized and graphed in EdgeR and Python, respectively. The light colored, shaded regions along the line graph represent the standard deviation at each time point.

### hfb9a is required for normal hydrophobicity and oxidative stress response

The gene encoding HFB9A was deleted via *Agrobacterium*-mediated transformation with a homologous recombination cassette (Supplementary Figure 1A). The knockouts were confirmed and screened for wild-type copies of the gene by PCR (Supplementary Figure 1B). The Δhfb9a mutants demonstrated a statistically significant difference in hyphal surface hydrophobicity as measured by contact angle of a water droplet on the surface of an agar plug (p < 0.05, Figure 4A). There was no significant difference in radial growth on PDA between the mutants and wild-type (Supplementary Figure 2). However, mutant strains grew significantly slower under oxidative stress than wild-type (p < 0.05, Figure 4B). The mutants grew similar as wild-type in confrontation with *P. ultimum* (Supplementary Figure 3A) and retained biocontrol activity against *R. solani* on cotton roots (Supplementary Figure 3B).

**Figure 4.**
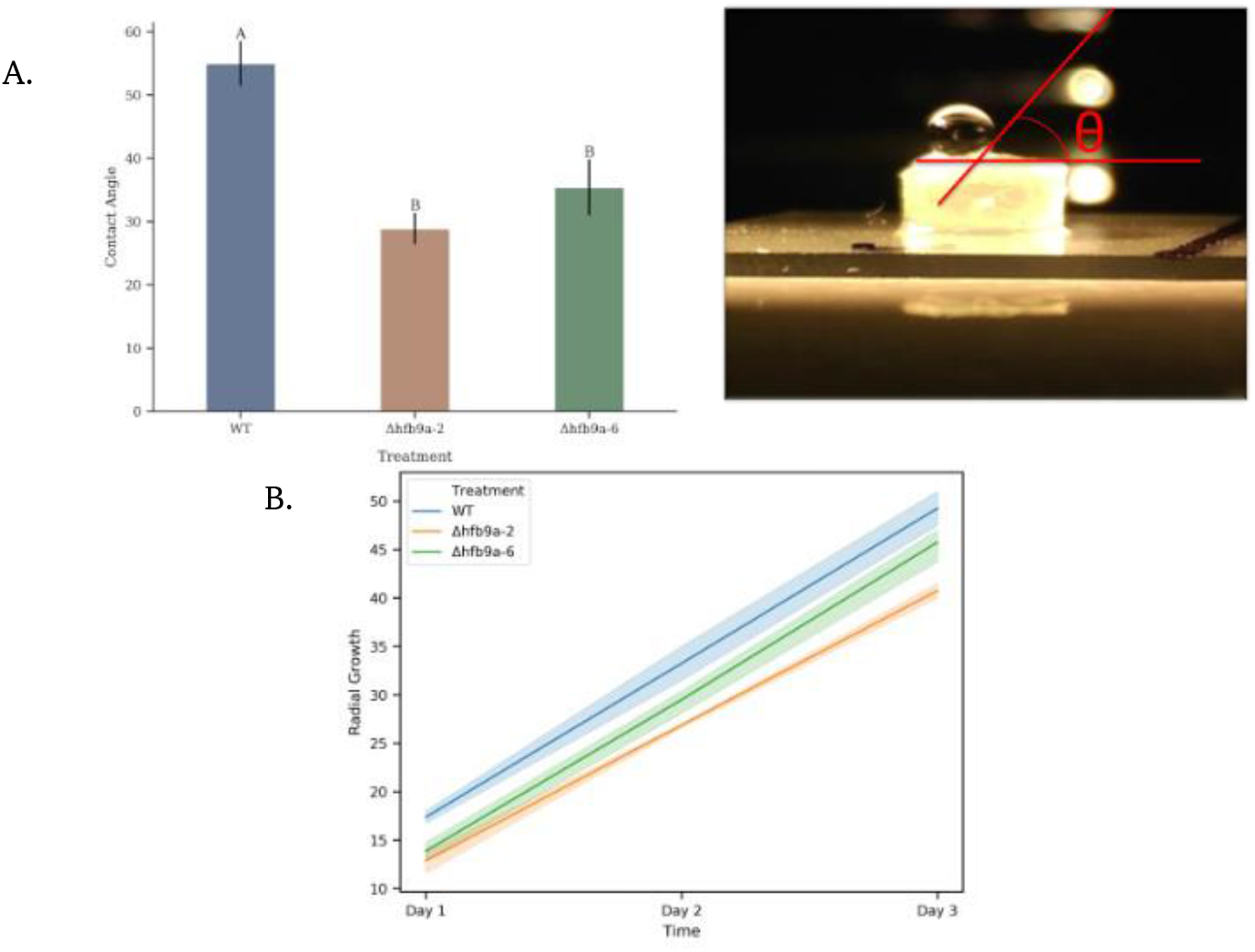
Surface hydrophobicity and oxidative stress response. **A.**Surface hydrophobicity of fungal mycelium as measured by contact angle of a water droplet as it lay on the surface of mycelium. Different letters represent statistically different groups (p < 0.05) as determined by ANOVA and Tukey’s HSD. Error bars indicate standard deviation. **B**. Radial growth of each strain over the course of three days on PDA amended with sodium menadione bisulfite to induce oxidative stress. Colored, shaded regions indicate standard deviation.

### hfb9a has a role in host root colonization and induction of systemic resistance

The ability of Δhfb9a mutants to colonize maize roots was significantly reduced in the hydroponic system (Figure 5A). Observations prior to harvest indicated a similar amount of each strain enveloping the roots. Interestingly, upon addition of pregerminated fungal tissue to the hydroponic medium, individual colonies could be seen attaching to the roots as the biomass became dispersed. This was significantly faster than expected and did not support our initial hypothesis that this hydrophobin was involved in attachment of the fungus to the roots. Additionally, visualization of the colonized roots by confocal microscopy indicated attachment to the root epidermis by the mutant, but only wild-type was able to colonize internal portions of the root sections extensively (Figure 5B, 5C).

**Figure 5.**
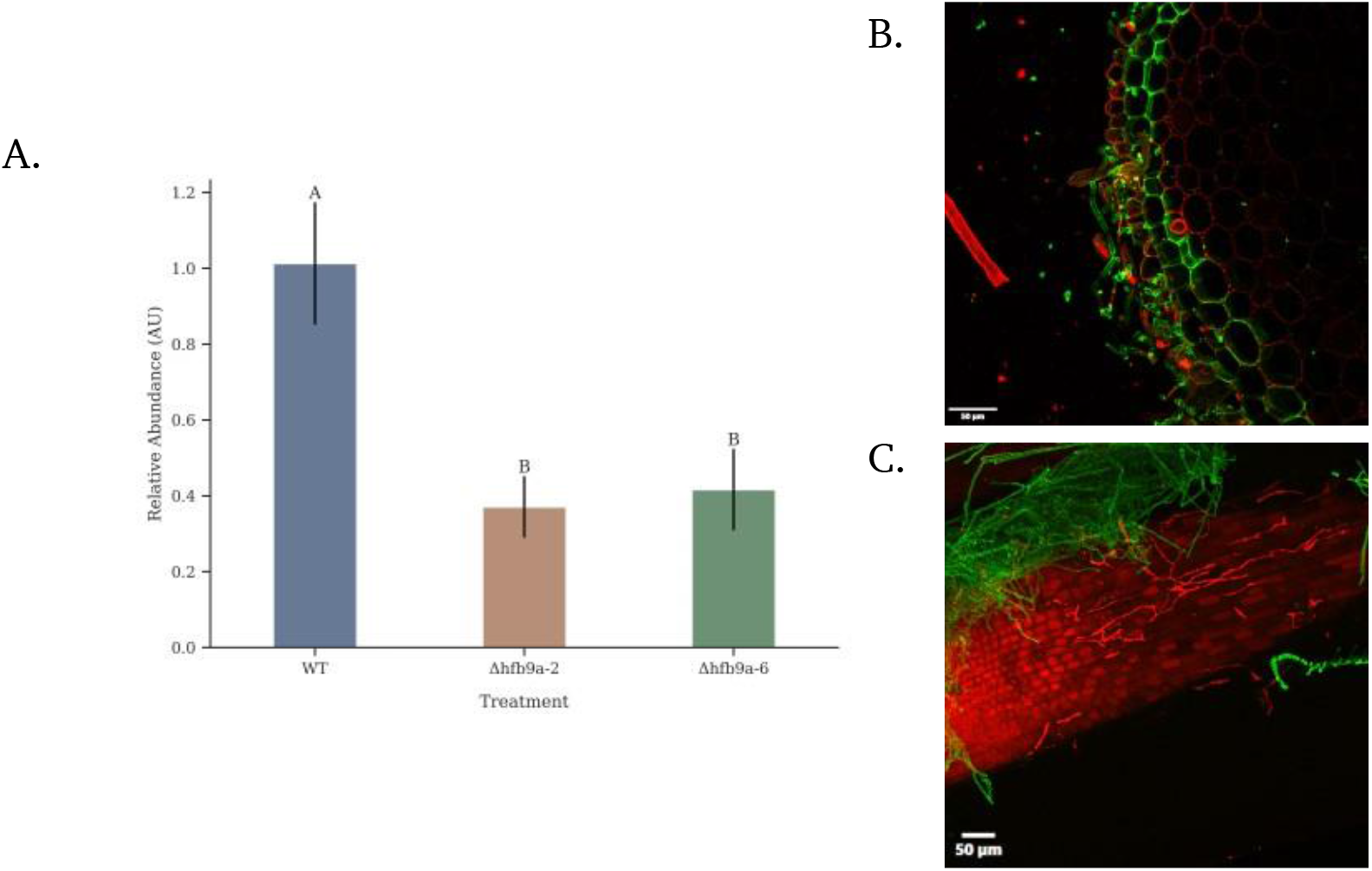
Root colonization of maize. **A.** Root colonization of maize roots in a hydroponic system by different strains of *T. virens.* Quantitative-PCR was used to determine the relative abundance of fungal DNA compared to maize DNA and normalized to the wild-type strain. A smaller number represents less colonization of maize roots by the fungus compared to the wild-type strain. Different letters represent statistically different groups (p < 0.05) as determined by ANOVA and Tukey’s HSD. Error bars indicate standard deviation. **B and C.** Confocal micrographs visualizing the wildtype strain **(B)** or Δhfb9a mutant strain **(C)** colonizing maize roots. Fungal tissue was stained with WGA-AlexaFluor-488 (green) and maize tissue was stained with propidium iodide (red). Individual hyphae can be seen in the intercellular spaces of the plant cells colonized by the wild-type strain. In contrast, the only observable hyphae present were extracellularly attached to the surface of roots colonized by Δhfb9a mutant strains.

The Δhfb9a mutants were analyzed for their ability to induce systemic resistance against *C. graminicola* in maize plants. The average lesion area of the plants treated with Δhfb9a mutants were significantly larger than the wild-type treated plants, indicating a lack of ISR (p < 0.05, Figure 6A, 6B). Plants treated with Δhfb9a mutants did not differ significantly in height or dry weight compared to wild-type treated plants (data not shown). Western blot analysis was performed to determine whether the reduction of ISR was due to decreased production of the elicitor protein SM1. Total protein was extracted from wild-type and Δhfb9a strains grown in PDB for 48 hr at 150 rpm. Each aliquot was diluted to 1 mg/ml and a western blot performed using anti-SM1 antibodies (Djonović*et al.*, 2006a). There was no appreciable difference in the amount of protein detected (Figure 7).

**Figure 6.**
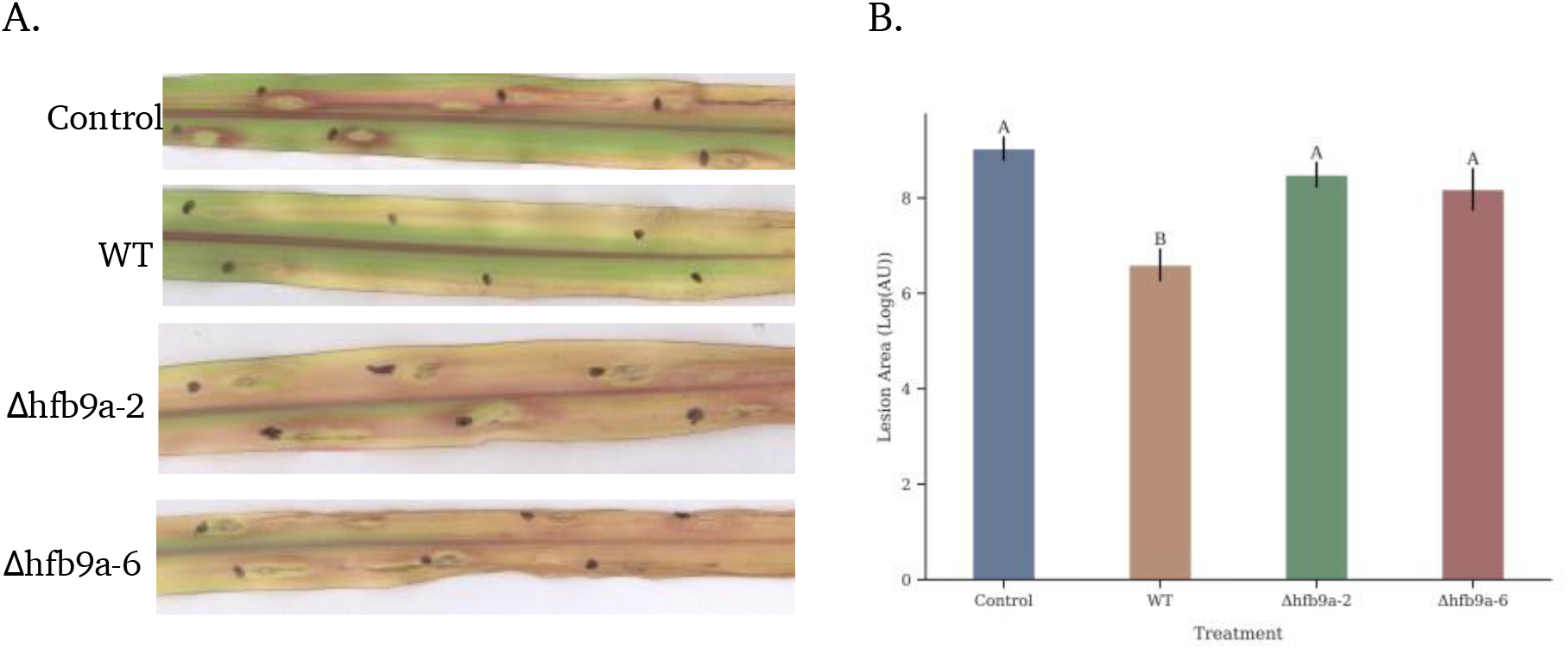
Induced systemic resistance of maize plants treated with the wild-type strain or Δhfb9a deletion mutants. Areas of individual lesions were measured in ImageJ. Different letters represent statistically different groups (p < 0.05) as determined by ANOVA and Tukey’s HSD. Error bars indicate standard deviation.

**Figure 7.**
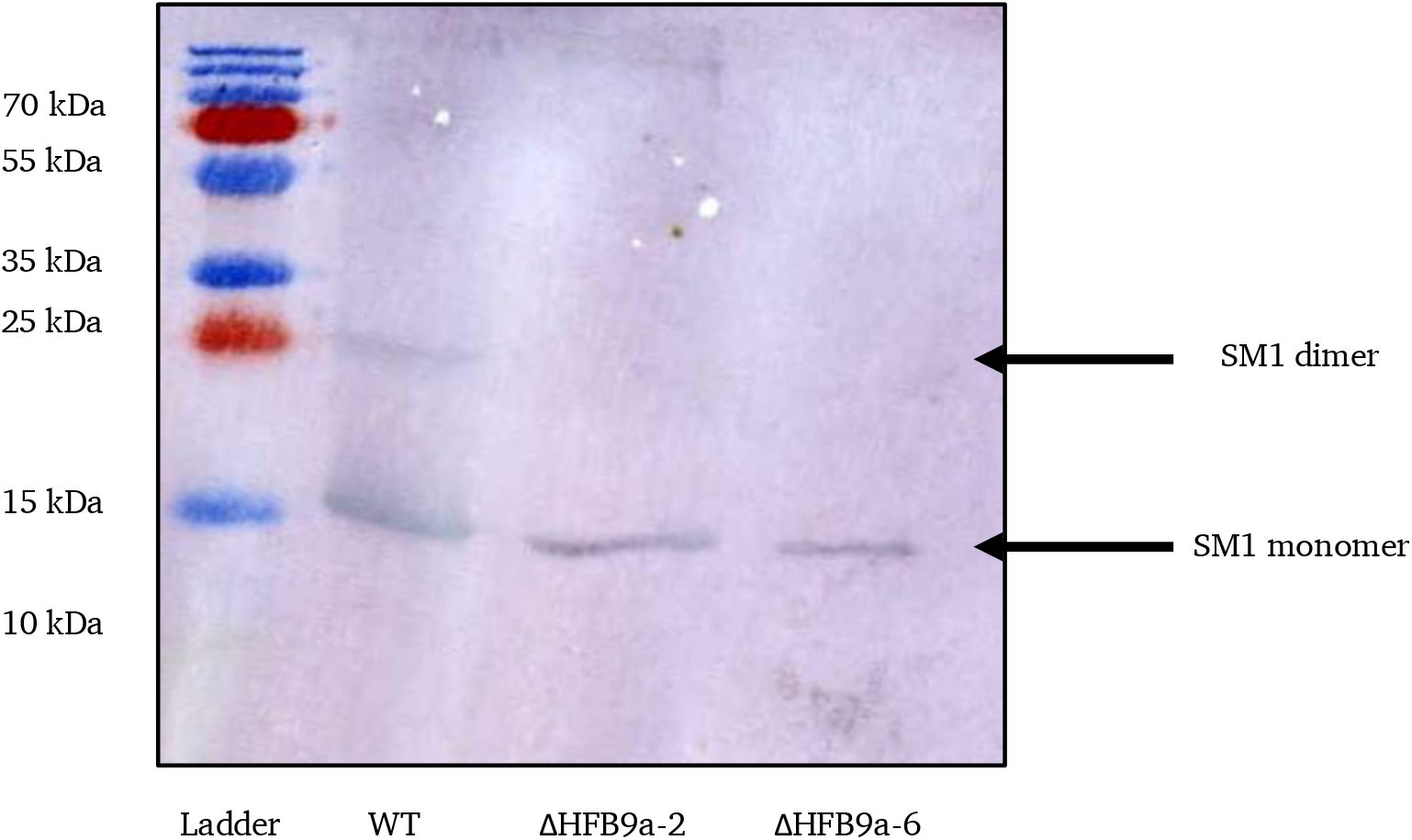
SM1 protein production. A western blot using antibodies specific to SM1 to determine the production of SM1 protein by the wild-type strain and deletion mutants (Δhfb9a-2 and Δhfb9a-6). All lanes were loaded with one ug of protein. There was no discernable difference in production of SM1 between the mutants and wild-type strains.

### hfb9a accelerates T. virens cellulase and chitinase activity

The Δhfb9a deletion mutants demonstrated approximately 90% less cellulase activity compared to the wild-type fungus (Figure 8A). To determine whether this effect was substrate specific, we duplicated the assay using colloidal chitin, pectin, and lignin. Chitinase activity on colloidal chitin was impacted by the loss of HFB9A (~95% reduction, Figure 8B). However, no impact on pectin or lignin degradation was found with this assay.

**Figure 8.**
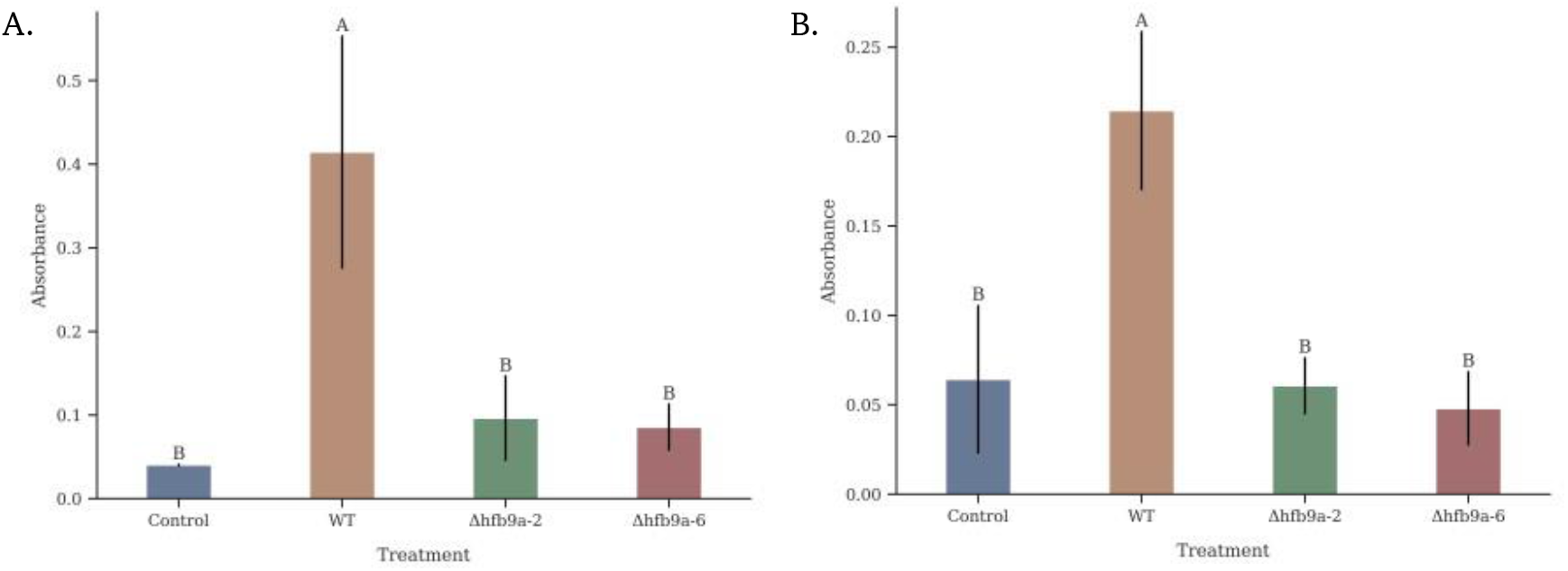
Enzyme activity determination. Cellulase (A) or chitinase (B) activity of culture filtrates from the wild-type strain and Δhfb9a deletion mutants as measured by the DNS assay. In each assay, the wild-type strain exhibited significantly higher enzymatic activity on cellulose and colloidal chitin (A and B, respectively). Different letters represent statistically different groups (p < 0.05) as determined by ANOVA and Tukey’s HSD. Error bars indicate standard deviation.

### HFB9A protein complements enzyme activity of mutants

The *E. coli* expression vector pET30B(+) was used to produce recombinant HFB9A with a fused 6xHis tag at the C-terminal end. Expression following induction with IPTG was attempted at 30°C overnight or 37°C for four hr. Protein production was only detected with cultures incubated at 37°C and present only in the insoluble fraction of the lysate.

The recombinant HFB9A protein was purified from solubilized inclusion bodies by nickel affinity chromatography and refolded on the column. Following elution with imidazole, the fractions containing the protein of interest were identified with a dot blot. The positive fractions were pooled, assayed by western blotting (Figure 9), and the protein was stored at 4°C for future use. The protein was collected in a buffer with high salt and imidazole. We attempted to exchange the buffer by dialysis but found that the protein aggregated and precipitated out of solution. To avoid aggregation and the resulting insolubility, the protein was rapidly precipitated using 4 volumes of ice-cold acetone; however, the protein remained insoluble upon the reconstitution attempts. The protein was re-solubilized via treatment with formic acid followed by the addition of an equal volume of 30% H2O2 to produce performic acid (Wosten *et al.*, 1993). The protein immediately went into solution following evaporation of the acid. The new protein solution retained the same surface activity in comparison with samples dissolved in the original buffer and was used throughout the rest of the study.

**Figure 9.**
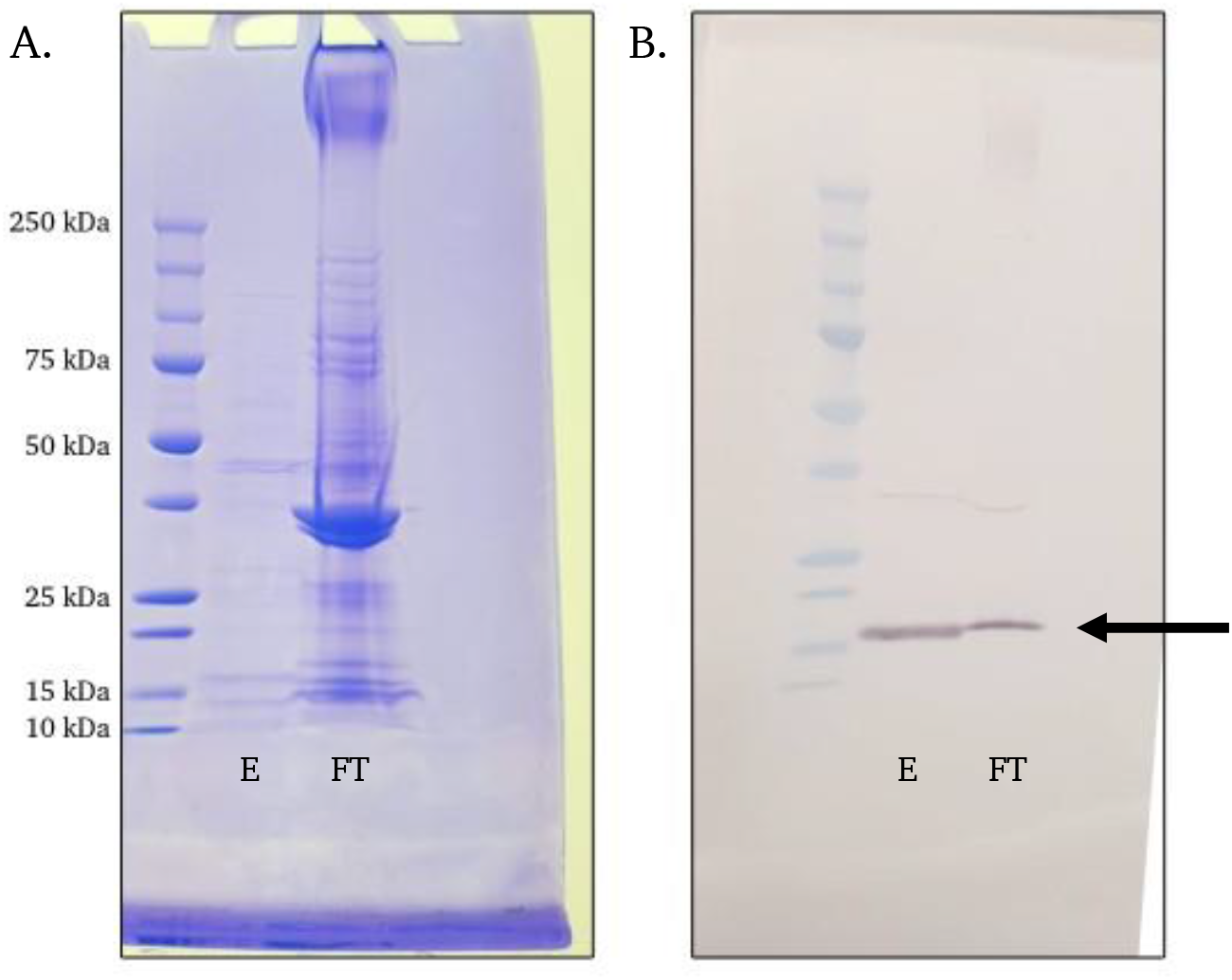
Detection of recombinant HFB9A. Coomassie Blue stained SDS-PAGE gel (**A**) and western blot (**B**) with antibodies specific to the H6 tag to detect recombinant HFB9A. E: eluted fraction, FT: column flowthrough prior to elution. The arrow indicates the band corresponding to recombinant HFB9A protein.

The purified protein was able to complement the cellulase activity of the mutants at 1 μM concentration. In addition to restoring the enzyme activity of the mutants to wild-type levels, addition of purified protein to the wild-type culture filtrate was able to enhance the cellulolytic activity against filter paper (~25% increase, Figure 10A). The HFB9A protein on its own displays no enzymatic activity and requires culture filtrate to have this effect. The protein buffer was also tested to ensure that there was no interaction in this assay and found no cellulolytic activity. Additionally, the pure protein at 1 μM concentration was able to enhance the activity of commercially sourced cellulase by 13% compared to the untreated control (Figure 10B).

**Figure 10.**
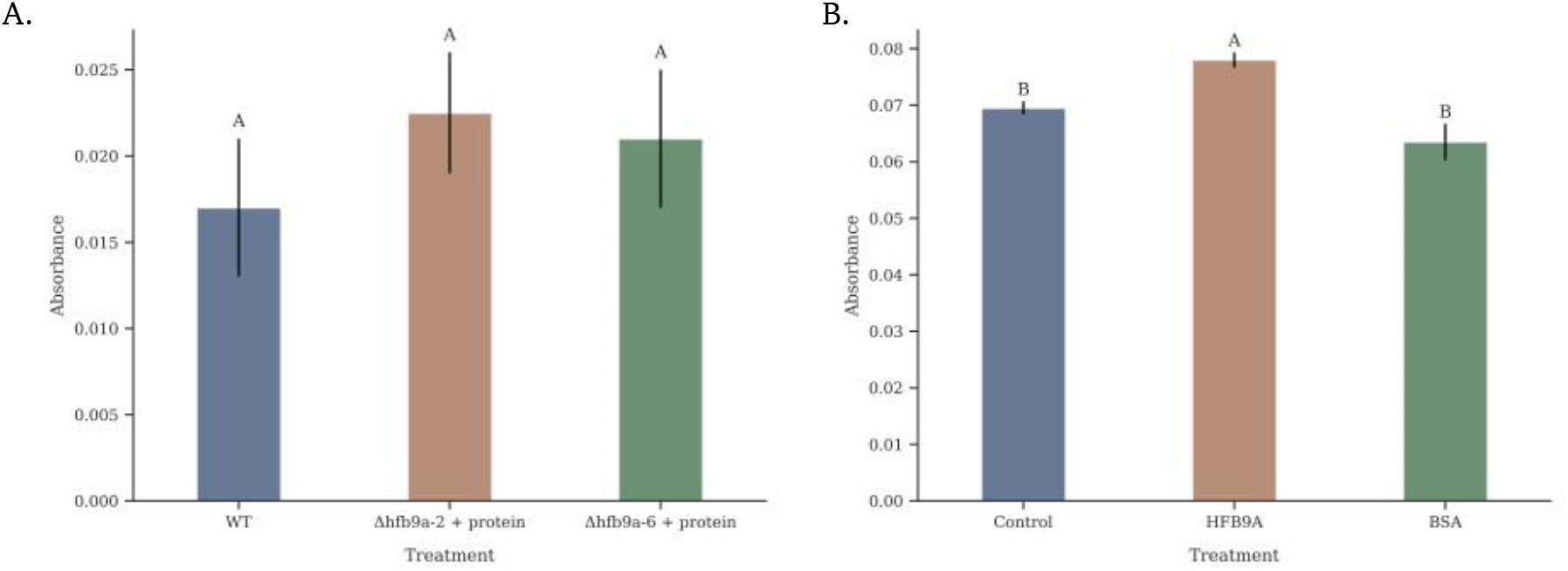
Enhancement of enzyme activity by recombinant HFB9A. **A.** Complementation of Δhfb9a deletion mutant cellulase activity by addition of purified HFB9A protein. Cellulase activity was measured by the DNS assay with a cellulose substrate. **B.** Enhancement of commercial cellulase with purified HFB9A protein. BSA treatment was included as a control at the same molar concentration as HFB9A. Different letters represent statistically different groups (p < 0.05) as determined by ANOVA and Tukey’s HSD. Error bars indicate standard deviation.

### hfb9a induces phosphorylation of AtMAPK 3 and 6

Purified protein samples of HFB9A and SM1 were applied to Arabidopsis seedlings to determine whether the proteins could induce rapid activation of plant innate immunity commonly associated with microbe associated molecular patterns (MAMPs) by phosphorylation of AtMAPK3 and AtMAPK6. HFB9A treated samples displayed phosphorylation of AtMAPK3 and AtMAPK6 starting 30-min post inoculation, whereas SM1 and buffer control samples did not display phosphorylation 15- or 30-min post inoculation (Figure 11).

**Figure 11.**
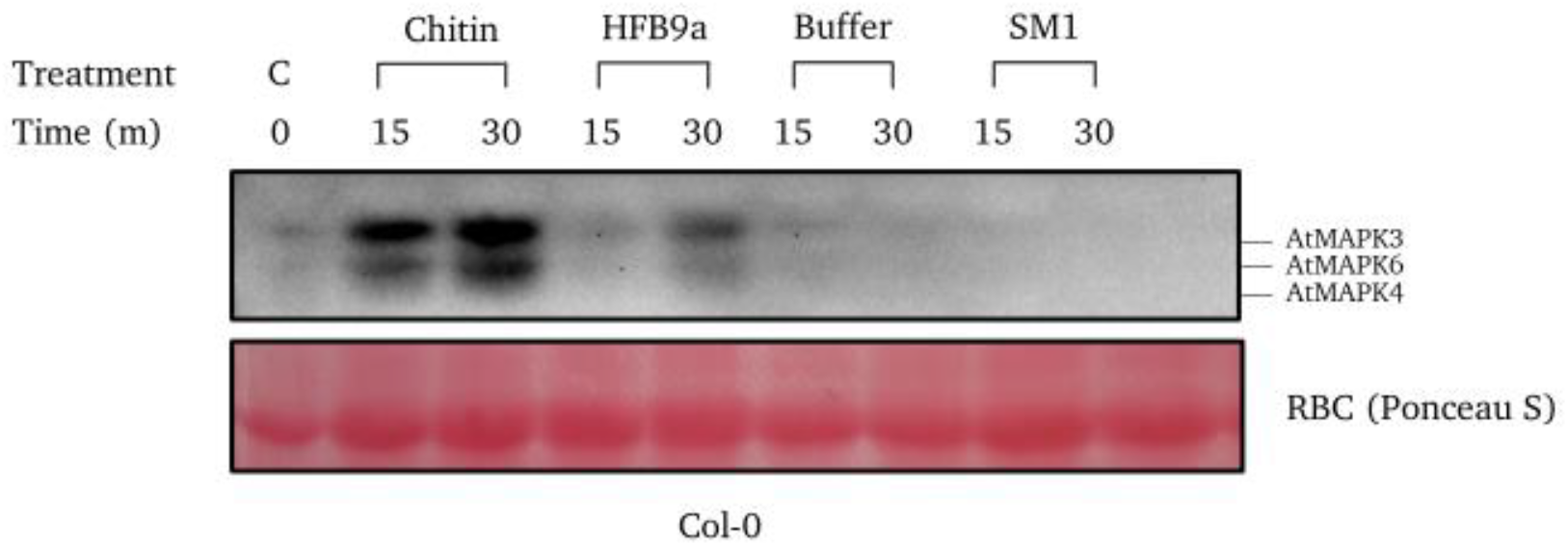
MAMP recognition by *Arabidopsis thaliana.* MAPK phosphorylation of *Arabidopsis* by addition of HFB9A over the course of 15- and 30-min. Phosphorylation was detected by antibodies specific to pERK1/2. Chitin was included as a positive control.

## Discussion

Hydrophobins are known to self-assemble at hydrophobic/hydrophilic interfaces. The resulting hydrophobin monolayer can modify the properties of the surface to which it adheres. Alteration of surface properties of materials by hydrophobins enable fungi to interact with previously intractable material as well as adapt to environmental changes. In this study, this phenomenon was demonstrated with HFB9A, as this hydrophobin is required for efficient colonization of maize roots and enhancement of cell wall degrading enzymes activity. Remarkably, ISR in maize treated with Δhfb9a deletion mutant strains was reduced, and in addition to reduced root colonization, one other potential mode of action of HFB9A on ISR may be due to the ability of the purified protein to induce activation of MAP kinase cascade, similarly to other MAMPs such as chitin.

It is not unprecedented that a hydrophobin may act as an elicitor of immunity to influence plant health. HYTLO1 was demonstrated to have a role in induced systemic resistance when applied to leaves, as well as when expressed transgenically in tomato plants (Ruocco *et al.*, 2015). The hydrophobin HFB9A may be functioning in a similar manner as a MAMP. AtMAPK3/6 proteins associated with MAMP response and innate immunity were phosphorylated after seedling treatment with purified HFB9A. The phosphorylation cascade starting with AtMAPK3/6 activates WRKY transcription factors involved in plant innate immunity (Adachi *et al.*, 2015). The direct activation of these proteins by purified HFB9A suggests that the fungal protein is recognized by the plant as non-self and induces defense responses. This result represents the first direct evidence of MAMP activity by a hydrophobin through induction of MAPK signaling cascades. Another protein with putative MAMP activity from *Trichoderma spp.* is SWOLLENIN from *T. asperellum*(Brotman *et al.*, 2008). Brotman *et al.* suggested that the carbohydrate binding domain of the protein acted as a MAMP, but they did not provide direct evidence, such as MAPK phosphorylation. Additionally, the production of SM1, a known elicitor of ISR, remained unchanged in the mutant strains in the presence of plant roots. However, there was an apparent difference in the dimerization of SM1 between the mutants and wild-type strains. The ability of SM1 to reduce disease progression by *C. graminicola* on maize leaves was shown to be affected by its dimerization state (Vargas *et al.*, 2008). Vargas *et al.* also hypothesized that glycosylation influenced the ability of SM1 to form dimers. It is tempting to speculate that HFB9a also influences aggregation of SM1, thus altering the plant response to the fungus. Remarkably, SM1 application to seedlings did not induce phosphorylation of MAPK proteins. This demonstrates, therefore, that the mechanism by which SM1 and HFB9A induce plant innate immunity is different at the molecular level. Further, it suggests that SM1 may not be perceived as a MAMP bu rather may act via as yet unknown mechanisms, potentially even as intracellular effector. SM1 and HYTLO1 have both been demonstrated to induce defense responses when applied to foliage (Djonović *et al.*, 2006b, Ruocco *et al.*, 2015). The lack of MAPK activation by SM1 suggests that HYTLO1 may induce defense more similarly to SM1 rather than act as a MAMP as previously hypothesized. The activation of MAPK signaling cascades by HFB9A, paired with decreased ISR as measured by foliar lesion size in plants treated with *HFB9a* deletion mutant strains, strongly suggests that HFB9A acts as a MAMP.

Our initial hypothesis was that HFB9A mediates attachment of hyphae to maize roots in a manner similar to TASHYD1 (Viterbo and Chet, 2006). However, based on observations of the attachment phenomenon in hydroponics systems and the timing of expression of the gene, this hypothesis is rejected. There is previous evidence that hydrophobins of both classes from *T. virens* enhance polyethylene terephthalate plastic degradation by cutinases (Przylucka *et al.*, 2014). The enzymatic enhancement of cellulases by HFB9A may aid the fungus in the degradation of plant cell walls, promoting colonization. The demonstrated reduction in cellulase enzyme activity and colonization by the Δhfb9a deletion mutants suggest this may be an additional role for some hydrophobins. The ability of HFB9A to enhance enzyme activity of cellulases and chitinases, but not other cell wall degrading enzymes such as pectinases may be due to the solubility of the substrates. Cellulose and chitin are both insoluble in water, whereas pectin is highly soluble. We speculate that HFB9A may be aiding the solubility of the substrates that the enzymes are acting upon, promoting enzyme attack. Beyond its biological role in the fungus, HFB9A could present a possibility for application in an industrial setting. Fungal hydrophobins are increasingly being used in commercial applications, such as in the protection of historic stonework from water damage (Winandy *et al.*, 2019). The cellulase and chitinase enhancement activity of HFB9A makes it a viable candidate for industrial applications, e.g. as an additive to enzyme cocktails.

Interestingly, the reduction in chitinase activity in Δhfb9a mutants did not influence the ability of the fungus to directly protect against *R. solani* infection of cotton roots, nor mycoparasitism of *P. ultimum* as measured by confrontation assays. It is possible that the minimal growth medium utilized to express cell wall degrading enzymes may not have been optimal for production of chitinases involved in mycoparasitism. A medium containing colloidal chitin or fungal cell walls might have been more appropriate for the production of mycoparasitism related chitinases.

Additionally, secondary metabolites such as gliotoxin and viridin have been demonstrated to have activity against both *R. solani* and *P. ultimum* (Howell *et al.*, 1993; Vargas *et al.*, 2014). The lowered activity of chitinases may be compensated by the activity of the antifungal secondary metabolites.

Overall, we demonstrate that HFB9A has an important role in the interaction between *T. virens* and its host. ISR is significantly reduced when HFB9A is not produced and the protein induces phosphorylation of MAPK proteins involved in immune responses, suggesting that HFB9A may function as a MAMP to activate ISR. Furthermore, the purified protein enhances the cell wall degrading activity of several cell wall degrading enzymes and has potential industrial applications.

## Acknowledgements

This work was funded by the Binational Science Foundation (Grant #2013202) awarded to CMK and BAH and USDA-NIFA (2016-67013-24730) awarded to CMK and MVK.

## References

Adachi, H., Nakano, T., Miyagawa, N., Ishihama, N., Yoshioka, M., Katou, Y., et al. (2015) Wrky transcription factors phosphorylated by mapk regulate a plant immune nadph oxidase in nicotiana benthamiana. Plant Cell27: 2645–2663.

Bayry, J., Aimanianda, V., Guijarro, J.I., Sunde, M., and Latgé, J.P. (2012) Hydrophobins-unique fungal proteins. PLoS Pathog 8: e1002700 http://dx.plos.org/10.1371/journal.ppat.1002700. Accessed July 29, 2016.

Bignell, E. (2012) The Molecular Basis of pH Sensing, Signaling, and Homeostasis in Fungi. In Advances in Applied Microbiology. Academic Press Inc., pp. 1–18.

Brotman, Y., Briff, E., Viterbo, A., and Chet, I. (2008) Role of swollenin, an expansin-like protein from Trichoderma, in plant root colonization. Plant Physiol 147: 779–789 http://www.pubmedcentral.nih.gov/articlerender.fcgi?artid=2409044&tool=pmcentrez&rendertype=abstract. Accessed November 13, 2015.

Correia, I., Prieto, D., Alonso-Monge, R., Pla, J., and Román, E. (2017) The MAP Kinase Network As the Nervous System of Fungi. Elsevier,.

Crutcher, F.K., Moran-Diez, M.E., Ding, S., Liu, J., Horwitz, B.A., Mukherjee, P.K., and Kenerley, C.M. (2015) A paralog of the proteinaceous elicitor SM1 is involved in colonization of maize roots by Trichoderma virens. Fungal Biol 119: 476–486.

Djonović, S., Pozo, M.J., Dangott, L.J., Howell, C.R., and Kenerley, C.M. (2006a) Sm1, a proteinaceous elicitor secreted by the biocontrol fungus Trichoderma virens induces plant defense responses and systemic resistance. Mol Plant-Microbe Interact 19: 838–853 http://apsjournals.apsnet.org/doi/abs/10.1094/MPMI-19-0838. Accessed August 31, 2016.

Djonović, S., Pozo, M.J., and Kenerley, C.M. (2006b) Tvbgn3, a β-1,6-glucanase from the biocontrol fungus Trichoderma virens, is involved in mycoparasitism and control of Pythium ultimum. Appl Environ Microbiol 72: 7661–7670.

Djonović, S., Vargas, W.A., Kolomiets, M. V., Horndeski, M., Wiest, A., and Kenerley, C.M. (2007) A proteinaceous elicitor Sm1 from the beneficial fungus Trichoderma virens is required for induced systemic resistance in Maize. Plant Physiol 145: 875–889 http://www.plantphysiol.org/cgi/doi/10.1104/pp.107.103689. Accessed October 4, 2016.

Guzmán-Guzmán, P., Alemán-Duarte, M.I., Delaye, L., Herrera-Estrella, A., and Olmedo-Monfil, V. (2017) Identification of effector-like proteins in Trichoderma spp. and role of a hydrophobin in the plant-fungus interaction and mycoparasitism. BMC Genet 18: 16 http://www.ncbi.nlm.nih.gov/pubmed/28201981. Accessed September 17, 2018.

Howell, C.R. (1987) Relevance of Mycoparasitism in the Biological Control of Rhizoctonia solani by Gliocladium virens. Phytopathology77: 992–994.

Howell, C.R., Stipanovic, R.D., and Lumsden, R.D. (1993) Antibiotic Production by Strains of Gliocladium virens and its Relation to the Biocontrol of Cotton Seedling Diseases. Biocontrol Sci Technol 3: 435–441 http://www.tandfonline.com/doi/abs/10.1080/09583159309355298. Accessed June 6, 2019.

Joensuu, J.J., Conley, A.J., Lienemann, M., Brandle, J.E., Linder, M.B., and Menassa, R. (2010) Hydrophobin fusions for high-level transient protein expression and purification in Nicotiana benthamiana. Plant Physiol 152: 622–33 http://www.ncbi.nlm.nih.gov/pubmed/20018596. Accessed March 31, 2017.

Kim, K.T., Jeon, J., Choi, J., Cheong, K., Song, H., Choi, G., et al. (2016) Kingdom-wide analysis of fungal small secreted proteins (SSPs) reveals their potential role in host association. Front Plant Sci 7.

Lamdan, N.L., Shalaby, S., Ziv, T., Kenerley, C.M., and Horwitz, B.A. (2015) Secretome of Trichoderma interacting with maize roots: Role in induced systemic resistance. Mol Cell Proteomics 14: 1054–1063 http://www.mcponline.org/content/14/4/1054.abstract?etoc.

Li, B., Jiang, S., Yu, X., Cheng, C., Chen, S., Cheng, Y., et al. (2015) Phosphorylation of trihelix transcriptional repressor ASR3 by MAP KINASE4 negatively regulates arabidopsis immunity. Plant Cell 27: 839–856.

Malinich, E.A., Wang, K., Mukherjee, P.K., Kolomiets, M., and Kenerley, C.M. (2019) Differential expression analysis of Trichoderma virens RNA reveals a dynamic transcriptome during colonization of Zea mays roots. BMC Genomics 20: 280 https://bmcgenomics.biomedcentral.com/articles/10.1186/s12864-019-5651-z. Accessed August 13, 2019.

Morán-Diez, M.E., Trushina, N., Lamdan, N.L., Rosenfelder, L., Mukherjee, P.K., Kenerley, C.M., and Horwitz, B.A. (2015) Host-specific transcriptomic pattern of during interaction with maize or tomato roots. BMC Genomics 16: 8 http://www.ncbi.nlm.nih.gov/pubmed/25608961. Accessed August 31, 2016.

Morris, V.K., Kwan, A.H., and Sunde, M. (2013) Analysis of the structure and conformational states of DewA gives insight into the assembly of the fungal hydrophobins. J Mol Biol 425: 244256 https://www.sciencedirect.com/science/article/pii/S0022283612008546?via%3Dihub. Accessed February 20, 2018.

Mustalahti, E., Saloheimo, M., and Joensuu, J.J. (2013) Intracellular protein production in Trichoderma reesei (Hypocrea jecorina) with hydrophobin fusion technology. N Biotechnol 30: 262–268.

Paz, Z., García-Pedrajas, M.D., Andrews, D.L., Klosterman, S.J., Baeza-Montañez, L., and Gold, S.E. (2011) One Step Construction of Agrobacterium-Recombination-ready-plasmids (OSCAR), an efficient and robust tool for ATMT based gene deletion construction in fungi. Fungal Genet Biol 48: 677–684.

Pieterse, C.M.J., Zamioudis, C., Berendsen, R.L., Weller, D.M., Wees, S.C.M. Van, and Bakker, P.A.H.M. (2014) Induced Systemic Resistance by Beneficial Microbes. Annu Rev Phytopathol 52: 347–375.

Przylucka, A., Ribitsch, D., Herrero-Acero, E., Gübitz, G., Kubicek, C.P., and Druzhinina, I. (2014) Hydrophobins class I versus class II: Trichoderma virens HFB9a and HFB9b (class I) are more effective as enhancing agents in enzymatic PET hydrolysis and surface modulators compared to HFB4 and HFB7 (class II). N Biotechnol 31: S190 https://www.sciencedirect.com/science/article/pii/S1871678414009972?via%3Dihub. Accessed August 17, 2018.

Ramírez-Valdespino, C.A., Casas-Flores, S., and Olmedo-Monfil, V. (2019) Trichoderma as a model to study effector-like molecules. Front Microbiol 10: 1030.

Ruocco, M., Lanzuise, S., Lombardi, N., Woo, S.L., Vinale, F., Marra, R., et al. (2015) Multiple Roles and Effects of a Novel Trichoderma Hydrophobin. Mol Plant-Microbe Interact 28: 167–179.

Saldajeno, M.G.B., Naznin, H.A., Elsharkawy, M.M., Shimizu, M., and Hyakumachi, M. (2014) Enhanced Resistance of Plants to Disease Using Trichoderma spp. In Biotechnology and Biology of Trichoderma. pp. 477–493.

Schneider, C.A., Rasband, W.S., and Eliceiri, K.W. (2012) NIH Image to ImageJ: 25 years of image analysis. Nat Methods 9: 671–675.

Seidl-Seiboth, V., Gruber, S., Sezerman, U., Schwecke, T., Albayrak, A., Neuhof, T., et al.(2011) Novel hydrophobins from trichoderma define a new hydrophobin subclass: Protein properties, evolution, regulation and processing. J Mol Evol 72: 339–351 http://link.springer.com/10.1007/s00239-011-9438-3. Accessed July 21, 2016.

Vargas, W.A., Djonović, S., Sukno, S.A., and Kenerley, C.M. (2008) Dimerization controls the activity of fungal elicitors that trigger systemic resistance in plants. J Biol Chem 283: 19804– 19815.

Vargas, W.A., Mukherjee, P.K., Laughlin, D., Wiest, A., Moran-Diez, M.E., and Kenerley, C.M. (2014) Role of gliotoxin in the symbiotic and pathogenic interactions of Trichoderma virens. Microbiol (United Kingdom) 160: 2319–2330.

Viterbo, A., and Chet, I. (2006) TasHyd1, a new hydrophobin gene from the biocontrol agent Trichoderma asperellum, is involved in plant root colonization. Mol Plant Pathol 7: 249–258 http://doi.wiley.com/10.1111/j.1364-3703.2006.00335.x. Accessed August 2, 2016.

Winandy, L., Schlebusch, O., and Fischer, R. (2019) Fungal hydrophobins render stones impermeable for water but keep them permeable for vapor. Sci Rep 9: 6264 http://www.nature.com/articles/s41598-019-42705-w. Accessed July 12, 2019.

Wösten, H. a (2001) Hydrophobins: multipurpose proteins. Annu Rev Microbiol 55: 625–646.

Wosten, H., Vries, O. De, and Wessels, J. (1993) Interfacial Self-Assembly of a Fungal Hydrophobin into a Hydrophobic Rodlet Layer. Plant Cell 5: 1567–1574 http://www.ncbi.nlm.nih.gov/pubmed/12271047. Accessed April 3, 2017.

Wösten, H.A.B., and Vocht, M.L. De (2000) Hydrophobins, the fungal coat unravelled. Biochim Biophys Acta - Rev Biomembr 1469: 79–86 http://www.ncbi.nlm.nih.gov/pubmed/10998570. Accessed January 12, 2017.

Yang, J., Yan, R., Roy, A., Xu, D., Poisson, J., and Zhang, Y. (2014) The I-TASSER suite: Protein structure and function prediction. Nat Methods 12: 7–8 http://www.nature.com/articles/nmeth.3213. Accessed April 4, 2018.

Zampieri, F., Wösten, H.A.B., and Scholtmeijer, K. (2010) Creating surface properties using a palette of hydrophobins. Materials (Basel) 3: 4607–4625.

