## Supplemental Figures for "A class I hydrophobin in *Trichoderma virens* influences plant-microbe interactions through enhancement of enzyme activity and MAMP recognition"

### Slide 1
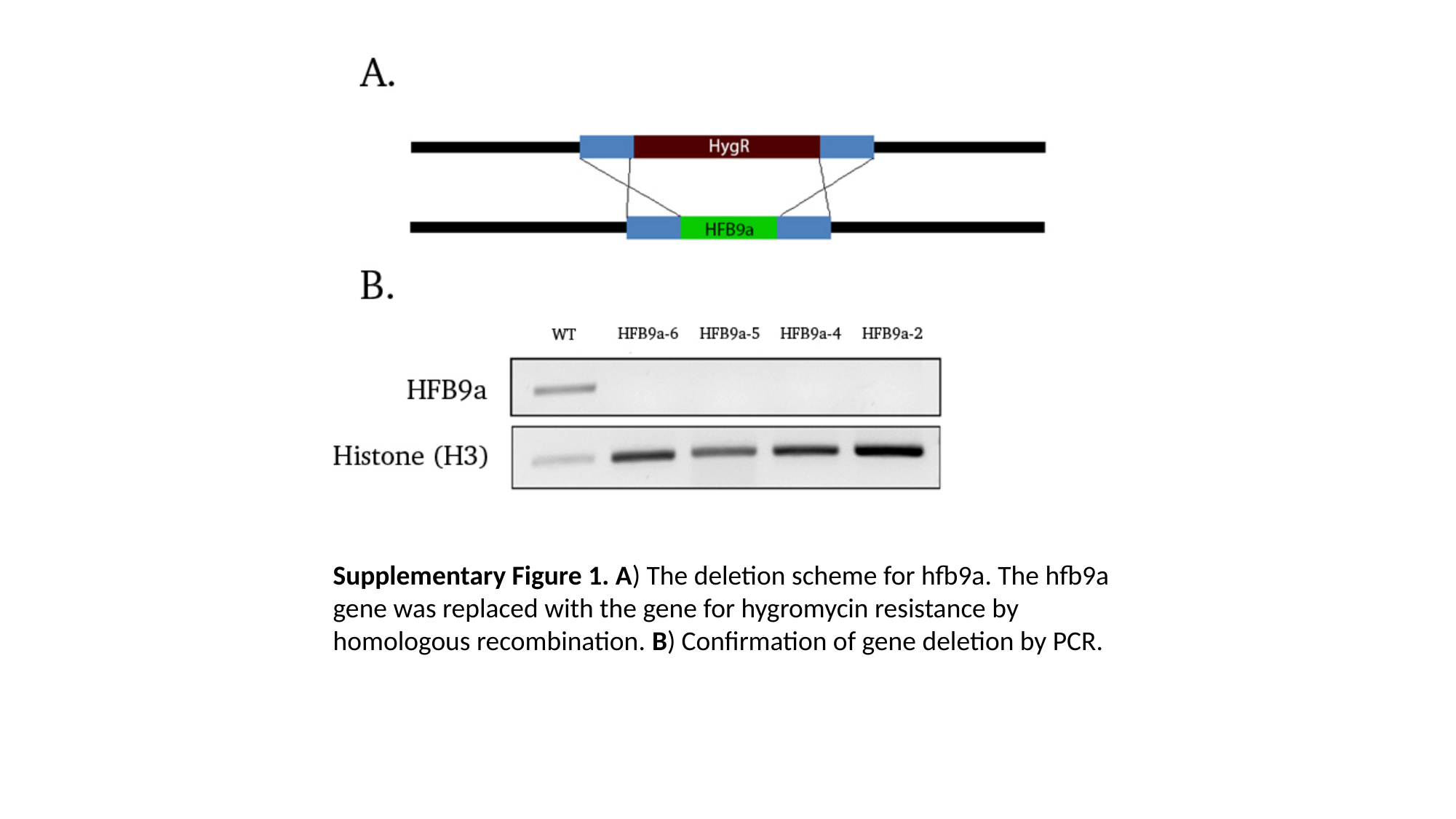

Supplementary Figure 1. A) The deletion scheme for hfb9a. The hfb9a gene was replaced with the gene for hygromycin resistance by homologous recombination. B) Confirmation of gene deletion by PCR.

### Slide 2
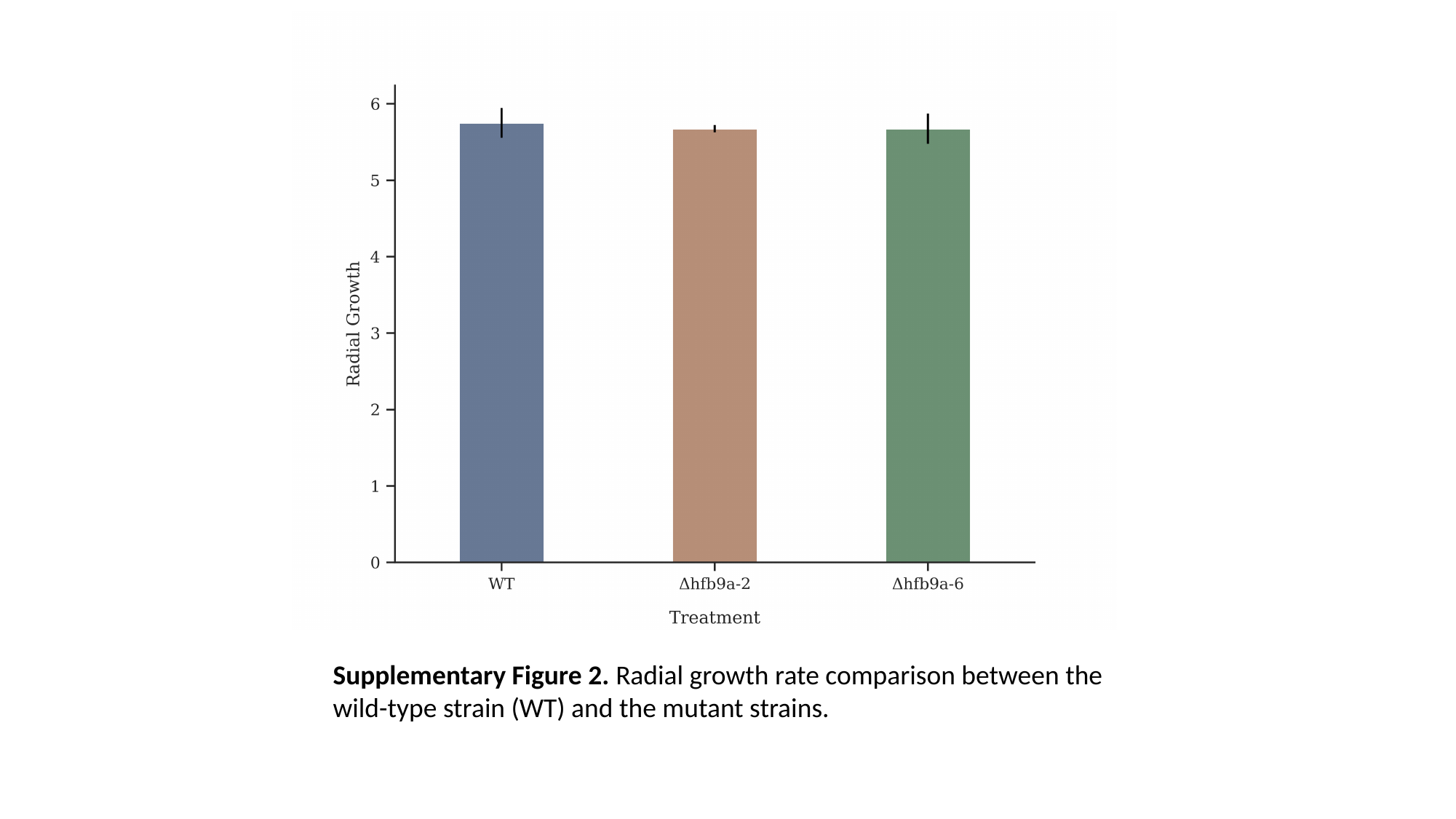

Supplementary Figure 2. Radial growth rate comparison between the wild-type strain (WT) and the mutant strains.

### Slide 3
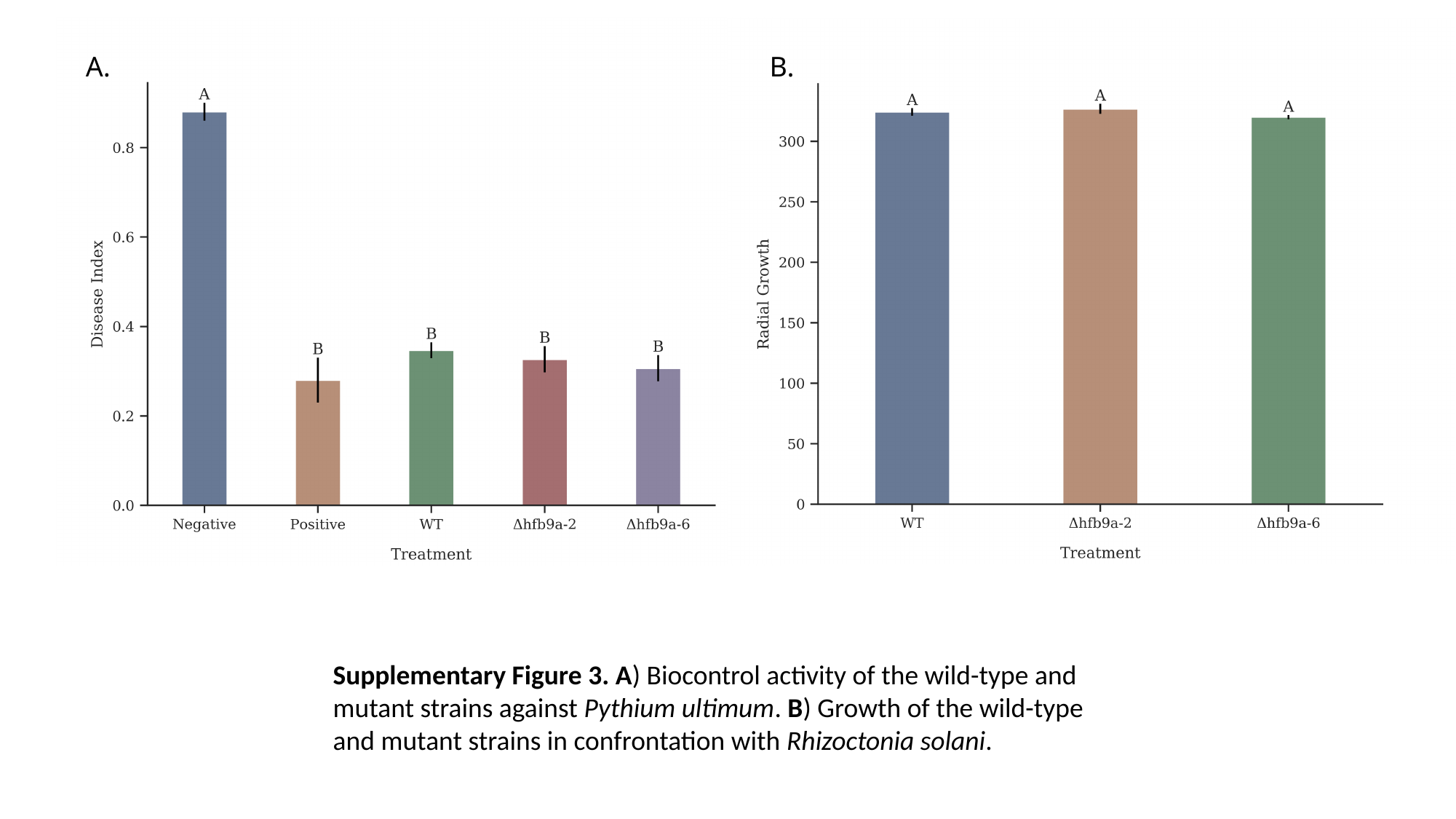

A.
B.
Supplementary Figure 3. A) Biocontrol activity of the wild-type and mutant strains against Pythium ultimum. B) Growth of the wild-type and mutant strains in confrontation with Rhizoctonia solani.
